# Molecular Landscape of Modality-Specific Exercise Adaptation in Human Skeletal Muscle through Large-Scale Multi-OMICs Integration

**DOI:** 10.1101/2024.07.14.603458

**Authors:** Macsue Jacques, Shanie Landen, Adam P Sharples, Andrew Garnham, Ralf Schittenhelm, Joel Stele, Aino Heikkinen, Elina Sillanpää, Miina Ollikainen, James Broatch, Navabeh Zarekookandeh, Ola Hanson, Ola Ekström, Olof Asplund, Séverine Lamon, Sarah E. Alexander, Cassandra Smith, Carlie Bauer, Mary N. Woessner, Itamar Levinger, Andrew E Teschendorff, Linn Gillberg, Ida Blom, Jørn Wulff Helge, Nicholas R Harvey, Larisa M Haupt, Lyn R Griffiths, Atul S. Deshmukh, Kirsi H Pietiläinen, Päivi Piirilä, Robert AE Seaborne, Bernadette Jones-Freeman, Nir Eynon

## Abstract

We conducted a large-scale, statistically powered, meta-analysis of exercise adaptations in human skeletal muscles, integrating epigenetic, transcriptomic, transcription factors, and proteomic data across 12 independent cohorts comprising over 1000 participants and 2340 human muscle samples. Our study identified distinctive signatures associated with maximal oxygen consumption (VO_2max_), and identified five genes robustly intersecting multi-OMIC layers. Notably, transcription factors predominantly functioned as activators across these layers, regulating expression of target genes irrespective of whether DNA methylation levels were low or high, indicating a synergistic effect between TFs and the methylome. Analysis of distinct exercise modalities (aerobic and resistance exercise) revealed unique gene pathways, contrasting with patterns observed in inactivity (muscle disuse) studies. These findings offer a comprehensive understanding of exercise and modality-specific adaptations, shedding light on muscle health and the molecular mechanisms associated with cardiorespiratory fitness, aging, and disease prevention.

## Introduction

Despite significant progress in understanding the effects of exercise on ageing and disease prevention, the molecular mechanisms underlying exercise adaptations remain unclear (Hoffman, 2017). A systematic approach that examines multi-OMIC layers within skeletal muscle, a key tissue for exercise adaptation, with sufficient statistical power, is essential to identify robust and novel exercise markers that can be targeted for therapeutic interventions and personalized exercise regimens.

Considerable attention has been directed towards understanding exercise adaptations in humans at the transcriptomic level to elucidate how gene expression changes with exercise. These studies have facilitated comprehensive meta-analyses (Pillon et al., 2020) and meta-regression analyses (Amar et al., 2021), identifying associations between the skeletal muscle transcriptome following both acute exercise and long-term training adaptations. Aerobic and resistance exercise training elicit some shared but predominantly distinct physiological and molecular responses. Aerobic training enhances oxidative metabolism and mitochondrial biogenesis, engaging genes closely associated with these pathways (Egan and Sharples, 2023). In contrast, resistance training stimulates predominantly muscle hypertrophy and strength gains, involving genes linked to muscle growth and remodelling (Amar et al., 2021; Turner et al., 2019). Pathways like AMP-activated protein kinase (AMPK) and Calcium/calmodulin-dependent protein kinase (CaMK) are crucial in aerobic training, promoting oxidative metabolism and mitochondrial biogenesis, whereas the mTOR pathway drives protein synthesis and muscle hypertrophy following resistance training (Qi et al., 2013). However, these studies often overlook significant layers of the multi-OMIC picture of exercise adaptation, including epigenetics and proteomic changes, which are critical to explore for a comprehensive understanding of the molecular mechanisms driving exercise-induced adaptations and their long-term health benefits. A recent study from the MoTrPAC consortium utilised this multi-OMIC approach in rat (MoTr et al., 2024), but this approach still needs to be applied in humans to develop a more comprehensive understanding of molecular adaptations to exercise training.

Epigenetics is the study of heritable changes in gene expression without alterations to the underlying DNA sequence. These changes affect gene activation or silencing and influence various biological processes and disease states (Lappalainen and Greally, 2017). Notably, epigenetic modifications are responsive to environmental stimuli, such as exercise (Jacques et al., 2019). DNA methylation, the most extensively studied epigenetic marker, has been associated with metabolism (Reid et al., 2017), numerous health conditions including cancer, type 2 diabetes, and metabolic syndrome (Crunkhorn, 2011; Heerboth et al., 2014; Kwak and Park, 2016), and ageing (Sharples et al., 2016; Voisin et al., 2021). DNA methylation alteration is associated with physiological adaptations post-exercise training (Howlett and McGee, 2016; Jacques et al., 2023a), with studies demonstrating genome-wide DNA methylation modifications in skeletal muscle with exercise training (Denham et al., 2015; Gorski et al., 2023; Lindholm et al., 2014; Nitert et al., 2012; Robinson et al., 2017; Ronn et al., 2013; Rowlands et al., 2014; Ruple et al., 2021; Seaborne et al., 2018a; Seaborne et al., 2018b; Turner et al., 2019). Human intervention studies are time consuming and invasive and therefore typically have small group sizes, these individual studies often lack sufficient statistical power due to the large number of CpGs investigated. For example, to achieve 95% power in typical exercise and DNA methylation studies (Jacques et al., 2019; Jacques et al., 2023; Voisin et al., 2024), a study would need approximately 316 tissue samples given effect sizes of 0.01-0.02 beta-value and standard errors average 0.06-0.07 beta-value. Therefore, a meta-analysis approach pooling multiple independent studies is ideal and cost-effective for drawing powerful and robust associations between DNA methylation and exercise adaptation.

Proteins, being the end products of the multi-OMIC layers, directly reflect molecular functionality (Molendijk et al., 2022). However, transcriptional changes do not always precisely indicate protein status due to the discordance in the timing of mRNA and protein expression. Additionally, numerous mechanisms independently regulate mRNA and protein abundance, including post-translational modifications, splicing, localization, and turnover (Koussounadis et al., 2015; Makhnovskii et al., 2020). Therefore, integrating the transcriptome with upstream (epigenome) and downstream (proteome) associations could provide a more comprehensive view of skeletal muscle adaptations to exercise training.

TFs play a significant role in regulating the transcription of genetic information from DNA to mRNA, similar to DNA methylation, by binding to specific DNA sequences known as transcription factor binding sites (TFBS) (Aguiari et al., 2021; Smith et al., 2020; Wardle, 2019). TFs are vital in orchestrating the molecular response to exercise training. Exercise-induced gene expression changes are often mediated by activation of a transcription factor, which binds to TFBS in the promoter regions of target genes, modulating their transcription. Key TFs involved in metabolic regulation, muscle growth, and adaptation include PPARδ, PGC-1α, and NF-κB (Egan and Sharples, 2023; Makhnovskii et al., 2020). Given that TFs typically bind to regulatory regions of the genome, exploring their relationship with DNA methylation is essential. This interaction is crucial because a specific TF might act as an activator while DNA methylation could serve as a repressor for the same gene, potentially attenuating or enhancing transcription depending on the interplay between these layers (Silva et al., 2022).

Here, we conduct a large-scale meta-analysis and integrate data from multiple OMICs sources. By harnessing a substantial sample size exceeding 1000 participants and 2340 human muscle samples, we aim to uncover robust and novel genes and pathways that may be overlooked by conventional single OMICs analyses. Furthermore, we aimed to uncover the connection between TFs, the methylome, and the transcriptome following exercise. We perform enrichment analyses of TFs and their binding sites, cross-referencing these results with our transcriptome findings, and examining their intricate interplay with DNA methylation patterns. Through our integrative approach, we construct a comprehensive functional correlation analysis across various OMICs layers (i.e. DNA methylation, transcriptome and proteome). Finally, to validate some of our discoveries related to exercise adaptation, we compare the exercise training outcomes with those from an inactivity (muscle disuse) meta-analysis (Deane et al., 2021) highlighting the divergent molecular changes induced by exercise compared with inactivity.

## Results

To clarify the study design and the results, we have subdivided our analysis into two parts. Part 1 focuses on OMICs analysis and its association with cardiorespiratory fitness (CRF) levels as assessed by VO_2max_. Part 2 examines OMICs analysis and changes after an exercise training intervention. Part 2 is further subdivided by exercise modalities (i.e. aerobic vs strength/resistance) where possible. **Supplementary Tables 1 & 2** and **Figure 1** provide summaries of the studies included in each part. Detailed descriptions of the analysis and included studies are in the ‘Methods’ section.

**Figure 1.**
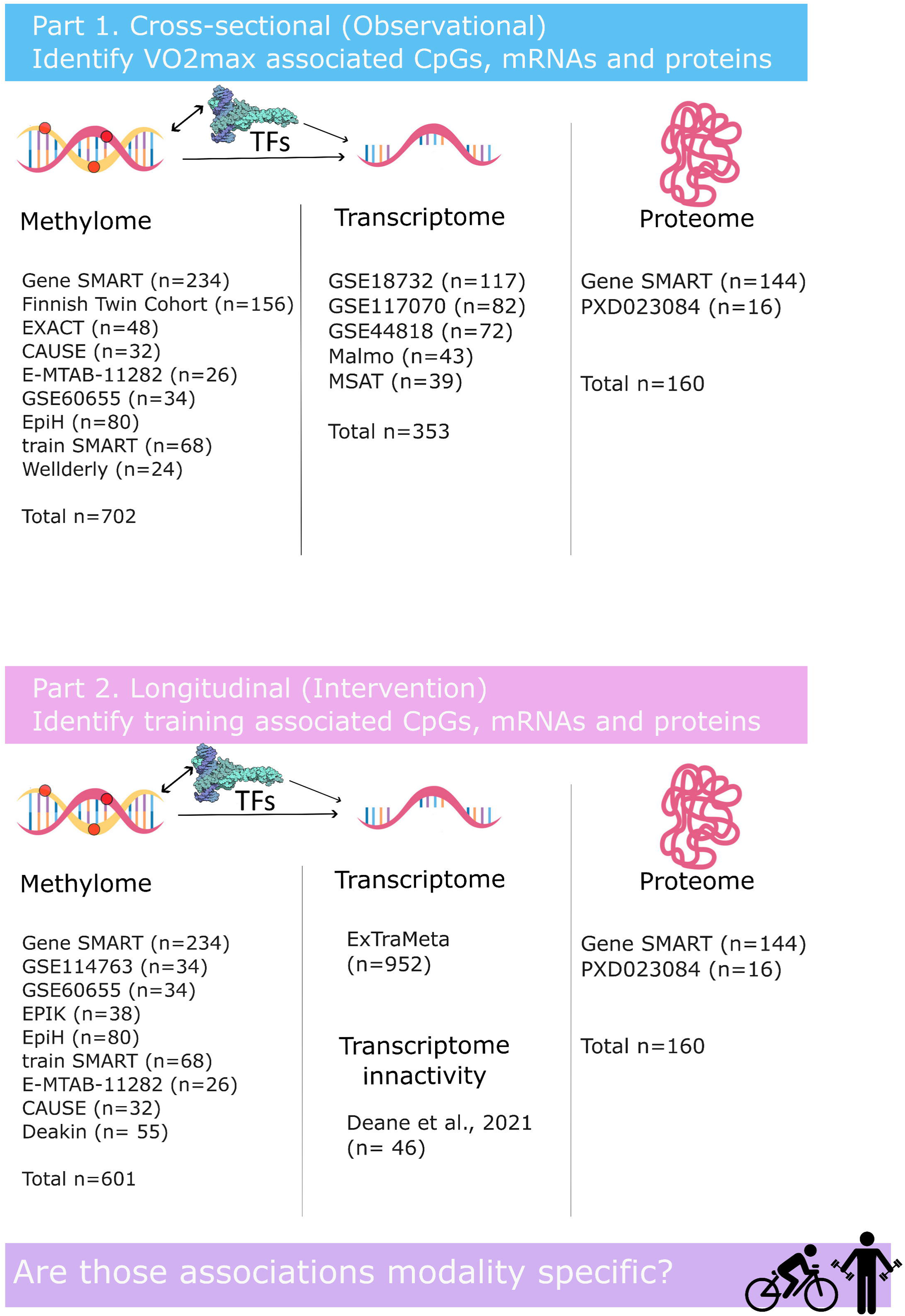
Study overview. First, we performed a large-scale data mining process to gather all existing omics from our lab, collaborators and open-access repositories (GEO). See ‘methods’ for inclusion criteria of each dataset and Supplementary Tables 1 & 2 for detailed information on each dataset. In Part 1, we identified cardiorespiratory fitness (measured as VO_2max_) associations with omics in human skeletal muscle by meta-analysing 702 samples across 9 cohorts for methylation, 353 samples across 4 cohorts for transcriptome and 160 samples across 2 cohorts for proteome. In part 2, we determined whether omics (methylation n=601 across 9 cohorts, transcriptome n=952 extracted from the ExTraMeta database and proteome n=160 across 2 cohorts) changed after exercise training and determined the modality-specific association in each of the OMIC layers when possible. In both parts we performed a series of OMIC integrations and pathway analyses to identify the molecular pathways affected by VO_2max_ and exercise training across OMIC layers. Finally, we have also investigated the intricate relationship between DNA methylation, TFs and transcription.

### Part 1 – Cardiorespiratory fitness omics signatures and integration

#### Methylome analysis depicts a large number CpGs associated with CRF to be mainly located in active chromatin states

We identified DNA methylation CpG sites associated with VO_2max_ across nine cohorts (n samples = 702). From a pool of 720,489 CpGs tested, and considering those examined in five or more studies to ensure robustness, we identified 9,822 CpG sites associated with VO_2max_ (FDR < 0.05, **Supplementary Table 3**). 43% of the CpGs exhibited a hypermethylated status correlated with higher VO_2max_, while 57% displayed a hypomethylation status associated with higher VO_2max_ (**Figure 2A**). Genes annotated to the top 10 CpG sites, including five hypo- and five hyper-methylated sites, are depicted in a forest plot (**Figure 2B**), with detailed information on the respective studies. This highlights the power of a meta-analysis approach: while top markers were generally not significant when considering individual studies (depicted by open circles in the forest plot), the combined meta-analysis reveals these targets as consistently and robustly changing across multiple studies, now achieving significance (depicted by the closed circles in orange). To avoid redundancy in our methylome findings, we performed regional analysis using *DMRcate*, identifying 742 differently methylated regions (DMRs) annotated to 983 genes associated with VO_2max_. Of these DMRs, 37% were hypermethylated, and 63% hypomethylated with higher VO_2max_ (**Supplementary Table 4**).

**Figure 2.**
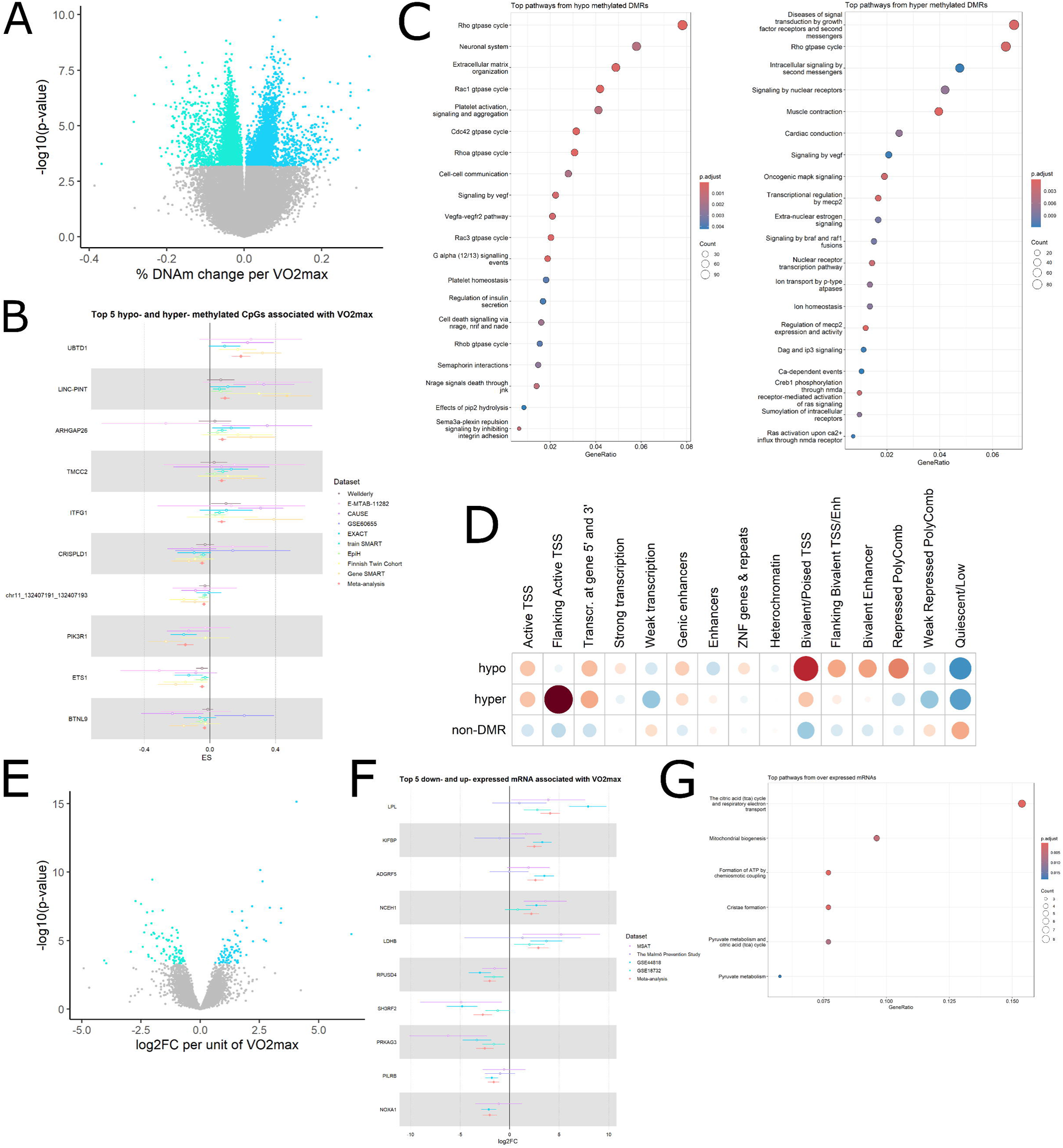
Methylome and Transcriptome meta-analysis reveals robust associations with CRF. **A)** Volcano plot of the effect size (% DNAm change per VO_2max_) and significance of VO_2max_ for each CpG included in the meta-analysis (pooled results of 9 independent muscle datasets); green coloured dots are CpGs that are differentially hypo-methylated and blue coloured dots are CpGs that are differentially hyper-methylated at FDR <0.05. **B)** Forest plots highlighting the top five hypo- (green) and hyper- (blue) methylated, showing estimates and confidence intervals in each dataset as well as the combined effect size observed in the meta-analysis. **C)** Overrepresentation analysis of the CRF associated Reactome pathways, subdivided based on hypo-or hyper-methylation CpGs. **D)** Distribution of chromatin states in male skeletal muscle from the Roadmap Epigenomics Project (Romanoski et al., 2015). The grids represent the residuals from the χ2-test, with the size of the circles being proportional to the cell’s contribution; red indicates an enrichment of the DMR category in the functional region, while blue indicates a depletion of the DMR category in the functional region. **E)** Volcano plot showing the effect size (log2FC per VO2max) and significance of VO_2max_ for each transcript in the meta-analysis (pooled results of 5 independent muscle datasets); green dots represent downregulated transcripts, while blue dots represent upregulated transcripts at FDR < 0.05. **F)** Forest plots highlighting the top five down- (green) and up- (blue) regulated transcripts, showing estimates and confidence intervals in each dataset as well as the combined effect size observed in the meta-analysis. **G)** Overrepresentation analysis of the CRF-associated Reactome pathways.

Gene expression can also be influenced by chromatin states, which are defined by specific combinations of histone modifications, methylation and DNA accessibility. Based on combinations of these epigenetic marks, 15 chromatin states have been described with distinct functional annotations (Romanoski et al., 2015). To further elucidate the association between the methylome and VO_2max_, we analysed the DMRs associated with VO_2max_ across chromatin states using the Roadmap Epigenomics database for skeletal muscle from males (Roadmap Epigenomics et al., 2015). This analysis revealed a non-uniform distribution across chromatin states (χ2-test p-value < 2.2 × 10e^−16^), with most DMRs associated with VO_2max_ being enriched in active chromatin states. Both hypo- and hyper-DMRs were under-represented in quiescent (silent) regions and weakly repressed PolyComb regions. Hypo-DMRs were strongly enriched in and around active TSS and enhancers, while hyper-DMRs were over-represented in Flanking Active TSS (**Figure 2D**).

Regarding genomic regulatory regions, 440 of the associated CpGs were situated in active transcriptional starting sites (TSS), and 3,082 were located in enhancer regions. Notably, the proportion of hypermethylation in these regions differed from the overall observed percentages for the significant CpGs (43% hyper-methylated and 57% hypo-methylated). In fact, when we overlapped our associated CpGs with the roadmap epigenomics database for skeletal muscle from males, we observed that 54% of our CpGs annotated to active TSS were hypermethylated, and 62% of CpGs annotated to enhancer regions also exhibited hypermethylation. Combined these findings suggest that gene bodies of skeletal muscle are more hypo-methylated and more variable versus more hyper-methylation in gene regulatory regions.

Our DMRs were over-represented in 133 Reactome pathways, including signalling by G protein coupled receptors (GPCR), GPCR ligand binding, circadian clock, antigen processing: ubiquitination & proteasome degradation, muscle contraction and mitochondrial biogenesis (**Figure 2C, Supplementary Table 5**).

#### Transcriptome analysis reveals multiple genes associated with CRF and a large number of enriched mitochondrial pathways

Previous transcriptome meta-analyses have primarily focused on exercise modality and other confounders such as sex and age. In our study, we conducted a meta-analysis of the transcriptome from 353 samples across five different studies, specifically investigating associations with CRF. We identified 162 transcripts significantly associated with VO_2max_ (FDR <0.05, **Supplementary Table 6, Figure 2E**). A forest plot exemplifying the top five upregulated and downregulated transcripts is presented in **Figure 2F**, detailing the studies where these targets were identified.

Overall, 50% of significant transcripts exhibited higher expression levels with higher VO_2max_ (**Figure 2E**). We identified ten Reactome pathways, all related to mitochondrial function, enriched among these transcripts (FDR <0.05, **Figure 2G, Supplementary Table 7**).

#### Methylome, transcription factors and transcriptome expression integration suggest synergistic interaction between OMICs with most enriched TFs acting as activators

To gain deeper insights into the mechanisms underlying CpGs associated with higher CRF, we conducted enrichment analysis of TF binding sites using UniBind (Puig et al., 2021). Hyper-DMRs were enriched for five unique TFs; Sine Oculis Homeobox Homolog 2 (*SIX2*), Peroxisome Proliferator-Activated Receptor Gamma (*PPARG*), Grainyhead Like Transcription Factor 2 (*GRHL2*), Neurogenin 2 (*NEUROG2*), and Estrogen Receptor 1 (*ESR1*) (FDR <0.05) (**Supplementary Table 8, Figure 3A**). Conversely, hypo-DMRs were enriched for 15 unique TFs, including Fli-1 Proto-Oncogene, ETS Transcription Factor (*FLI1*), ETS Proto-Oncogene 1, Transcription Factor (*ETS1*), ETS-Related Gene (*ERG*), RELA Proto-Oncogene, NF-KB Subunit (*RELA*), Nuclear Factor Erythroid 2-Related Factor 2 (encoded by NFE2L2) (*NRF2*), Runt Related Transcription Factor 1 (*RUNX1*), Spi-1 Proto-Oncogene, ETS Transcription Factor (*SPI1*), GATA Binding Protein 2 (*GATA2*), Nuclear Receptor Subfamily 3 Group C Member 1 (Glucocorticoid Receptor) (*NR3C1*), E74 Like ETS Transcription Factor 1 (*ELF1*), MYC Proto-Oncogene, BHLH Transcription Factor (*MYC*), MYCN Proto-Oncogene, BHLH Transcription Factor (*MYCN*), Jun Proto-Oncogene, AP-1 Transcription Factor Subunit (*JUN*), ETS Variant 6 (*ETV6*), BCL6 Transcription Repressor (*BCL6*) (FDR <0.05) (**Supplementary Table 8, Figure 3B**).

**Figure 3.**
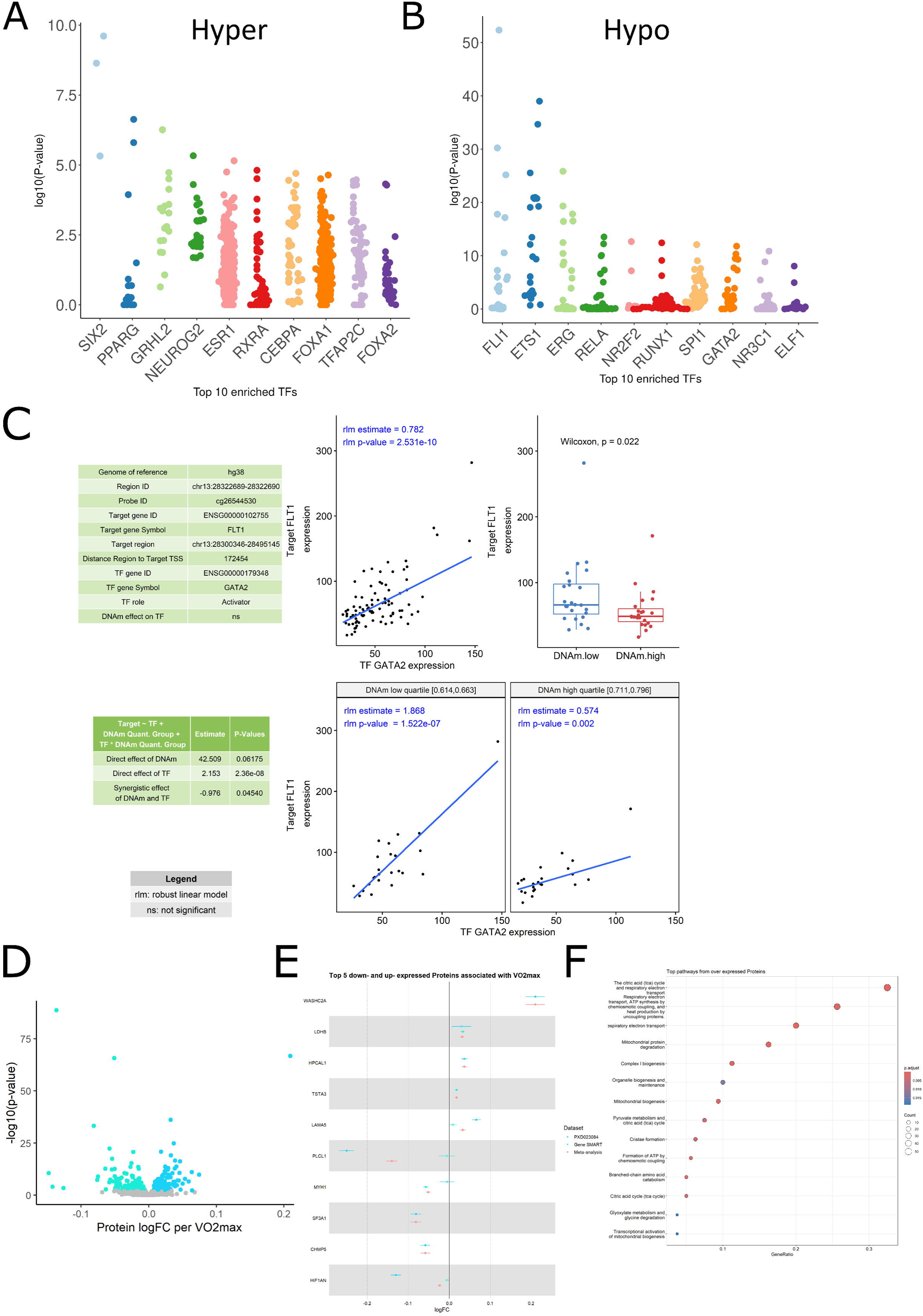
Integration analysis between DNA methylation, TFs and transcriptome. **A)** Top 10 transcription factors derived from hyper-DMRs. **B)** Top 10 transcription factors derived from hypo-DMRs. A & B were generated by the UniBind tool and represent enriched TFs before FDR correction. **C)** Output from MethReg analysis for the matched samples. The first table shows the triplet (CpG, TF, target gene) metadata, TF role (activator) and DNA methylation effect on the TF (ns). The second table shows the results from fitting robust linear model target gene expression residual ∼ DNAm group + TF + TF × DNAm group, where DNAm group is 0 if the sample has low DNA methylation levels (in the lowest quartile) at the given CpG or 1 if the sample has high methylation levels (in the highest quartile), indicating non-significant DNAm × TF interaction effect (P < 0.06). Top scatter-plot represents all samples are considered, there is significant association between target gene expression and TF activity (each dot represents a sample). Boxplot represents the comparison of target gene expressions between the DNAm groups shows samples with lower DNA methylation have higher target gene expression. Bottom scatter plots of target gene expression versus TF activity stratified by DNAm group (only samples in DNAm high or low groups are shown). In samples with low DNA methylation, TF activates target gene expression. In samples with high DNA methylation, target gene expression is still activated by TF in an attenuated manner. Therefore, TF is predicted to be an activator and DNAm is predicted to attenuate the effect of TF on the target gene. Abbreviations: DNAm, DNA methylation. **D)** Volcano plot of the effect size (log2FC per VO2max) and significance of VO2max for each protein included in the meta-analysis (pooled results of 5 independent muscle datasets); green dots represent proteins that are downregulated and blue dots represent proteins that are upregulated at FDR < 0.05. **E)** Forest plots highlighting the top five down- (green) and up- (blue) regulated proteins, showing estimates and confidence intervals in each dataset as well as the combined effect size observed in the meta-analysis. **F)** Overrepresentation analysis of the CRF associated Reactome pathways.

To further investigate the connection between significant TFs and the genome, we examined the genomic regions where these TFs are likely to bind. TFs enriched in hyper-DMRs collectively exhibited 59,508 transcription factor binding sites (TFBS) across both coding and non-coding regions of the genome. Within this group, 38,262 TFBS were situated in regions corresponding to known genes, as detailed in **Supplementary Table 9**. TFs enriched in hypo-DMRs collectively showed 59,318 TFBS, with 38,212 situated within known genes.

Of the 162 significant transcripts associated with VO_2max_, 157 were derived from genomic regions containing at least one TFBS for the top 20 enriched TFs derived from our hyper- and hypo-DMRs (**Supplementary Table 10**). A hypergeometric test confirmed that the observed overlap between differential DNA methylation and the enrichment for TFs, as well as between TFBS and transcriptome association targets, are highly unlikely to be explained by chance alone (p-value = 0.00031).

Gene expression can be regulated by various mechanisms, including DNA methylation and TF binding. Using MethReg, we performed an integrative analysis combining matching DNA methylation and gene expression (n=89 samples) with TFBS data from JASPAR2020 (Castro-Mondragon et al., 2022). This analysis revealed 79 significant interactions between methylome and TFs modulating gene expression for CpGs located in promoter regions. When considering a window of 500kb around the CpGs, we identified 1,941 significant interactions (DNAm:TF FDR <0.05) and 1,067 significant interactions when considering the five nearest upstream and downstream genes (**Supplementary Tables 11 to 13**).

In the context of CRF, the most significantly enriched TFs act as activators highly associated with target gene expression, with no significant effect of DNA methylation on TF activity. For example, the TF GATA2 showed a significant association with the target gene FLT1 (p-value = 2.5e-10), inferring the role of GATA2 as an activator of FLT1 regardless of DNA methylation level. This is corroborated by the highly significant direct effect of the TF (p-value = 2.36e-08) compared to the non-significant effect of DNA methylation (p-value = 0.06). In this case, low DNA methylation levels correlate with increased gene expression due to abundant TF binding (**Figures 3C**). However, high DNA methylation levels do not completely repress FLT1 expression possibly indicating reduced TF binding due to methylation. This example illustrates a synergistic interaction between DNA methylation and TFs in regulating target gene expression.

#### Proteome analysis depicts similar enriched pathways to those observed in the transcriptome analysis

We identified 351 proteins associated with VO_2max_ in the Gene SMART and PXD023084 cohorts (n=160) (FDR <0.05, **Supplementary Table 14**), with 58.1% of these proteins showing higher expression levels correlated with higher VO_2max_ (FDR<0.05, **Figure 3D**). A forest plot depicting five of the top up- and down-regulated proteins is provided in **Figure 3E**. Downstream analysis revealed significant enrichment in 18 Reactome pathways at the proteome level, with most pathways related to mitochondrial function, energy production and metabolism (FDR <0.05, **Figure 3F, Supplementary Table 15**).

#### Methylome, transcriptome and proteome integration reveal five intersected genes across OMICs and novel pathways associated with CRF

We identified five genes associated with VO_2max_ across all three OMIC layers; Neutral cholesterol ester hydrolase 1 (*NCEH1*), Aldehyde dehydrogenase 6 family, member A1 (*ALDH6A1*), Heat shock protein Family A Member 2 (*HSPA2*), Parkinsonism Associated Deglycase (*PARK7*) and Calcium Binding Protein 39 (*CAB39*). A hypergeometric test indicated that the overlap observed between significant differentially methylated genes (DMGs) differentially expressed genes (DEGs) and differentially expressed proteins (DEPs) cannot be attributed to chance alone (p-value=0.000625, **Figure 4A**).

**Figure 4.**
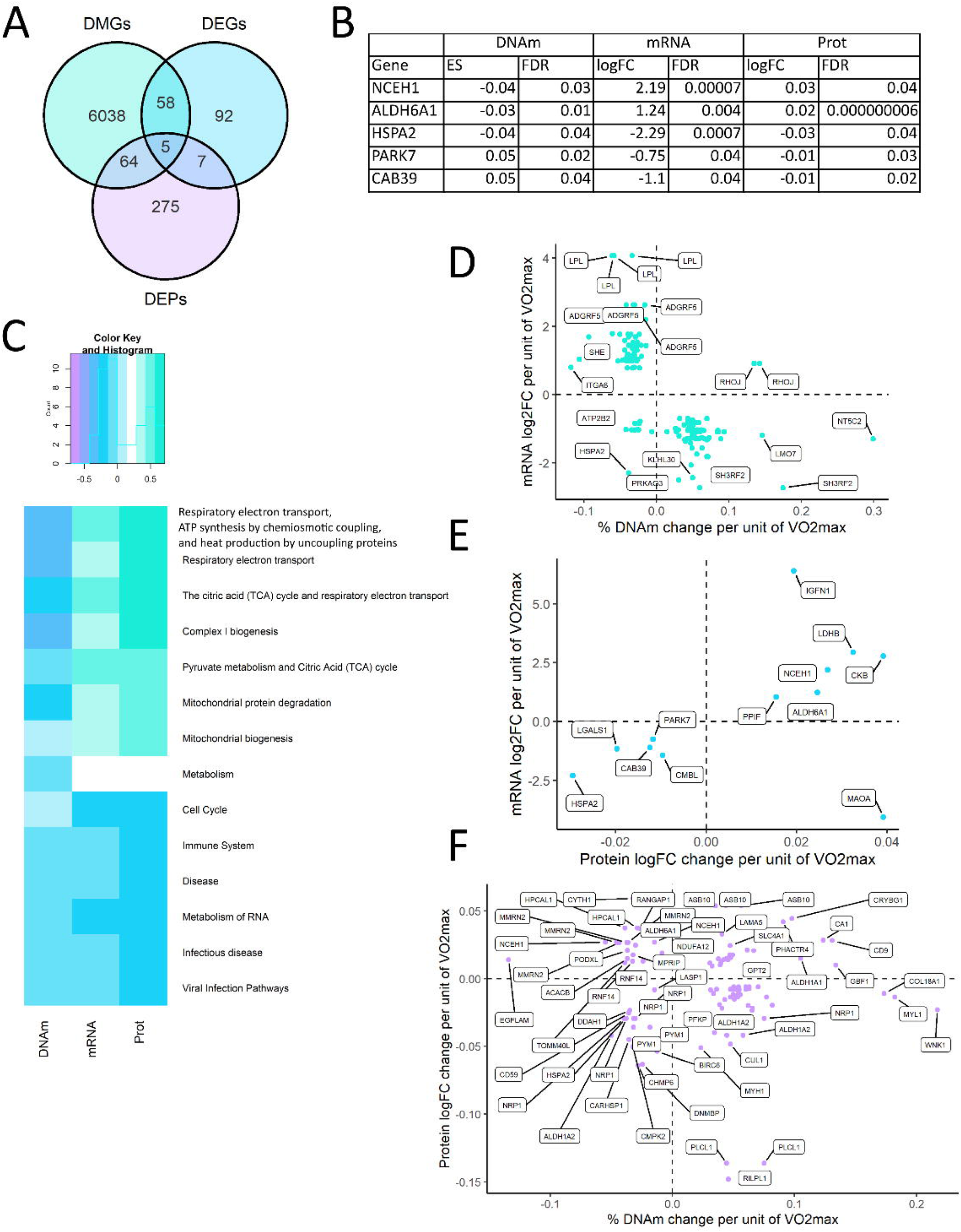
Overlap between VO2max differentially methylated genes (DMGs), differentially expressed genes (DEGs) and differentially expressed proteins (DEPs). **A)** Venn diagram representing targets overlap between omics. We identified 6,165 DMGs genes, 63 of which were also DEGs, and 69 were also DEPs. **B)** Genes associated with VO_2max_ that are intersected genes in all OMIC layers. **C)** Top integration pathways representing a clear inverse relationship between DNAm and mRNA as well as proteome. The blue colour represents an overall decrease in genes extracted from DNA methylation sites measured by the S distance, or mRNA/Protein expression belonging to the same pathway, and the green colour represents an increase in genes extracted from DNA methylation sites, or mRNA/Protein expression belonging to the same pathway. The band is represented in arbitrary units. **D)** OMIC pairs DNAm vs mRNA per unit of VO_2max_ **E)** Proteins vs mRNA per unit of VO_2max_ **F)** DNAm vs proteins per unit of VO_2max_. Each dot represents an intersected significant gene in both layers. Note that some genes in graphs D & F appear multiple times on the plot as multiple DMPs could be annotated to the same gene (e.g. LPL, RHOJ).

*PARK7* and *CAB39* exhibited increased mean methylation across CpGs with a concomitant decrease in gene expression and protein levels in association with higher VO_2max_. Conversely, *NCEH1* and *ALDH6A1* showed a decreased mean methylation with a concomitant increase in mRNA expression and protein abundance, with higher VO_2max_. Interestingly, *HSPA2* displayed an inverse relationship between DNA methylation and both gene expression and protein abundance (**Figure 4B**). Across all three OMIC layers *HSPA2* exhibited decreased effect size for DNA methylation, gene expression and protein abundance with increased VO_2max_.

To further elucidate these relationships, we investigated the chromatin state locations of significant CpGs for *HSPA2* according to the Roadmap Epigenomics database (Roadmap Epigenomics et al., 2015). In males, the CpG sites were located in the promoter region of *HSPA2* (termed “active transcript starting site”), while in females, it was located in enhancer regions. The methylome presented an inverse correlation with most intersected mRNA targets (**Figure 4D**), while the methylome and proteome presented a less clear directional relationship, with a mix of direct and indirect changes (**Figure 4F**). The mRNA and protein levels presented a nearly perfect relationship for the intersected targets, with only one exception Monoamine oxidase A (MAOA) showing decreased mRNA expression alongside increased protein expression (**Figure 4E**).

To identify pathways affected across all three OMIC levels, we performed a multi-contrast gene set enrichment analysis using a rank-MANOVA approach (Kaspi and Ziemann, 2020). We identified 210 Reactome pathways associated with VO_2max_ (**Supplementary Table 16**), with glyoxylate metabolism and glycine degradation showing the largest effect size (p-adjust MANOVA=0.0014, effects sizes measured by S ranking distance), and citric acid (TCA) cycle and respiratory electron transport showing the most significant associations (**Figure 4C**). Overall, pathways derived from the DNA methylation signatures of VO_2max_ were negatively correlated with mRNA expression and the protein level signature pathways (**Figure 4C**), while mRNA expression and protein level signatures were positively correlated with each other. The integration revealed an inverse relationship at the DNA methylation, mRNA, and protein level for pathways related to mitochondrial function, where a decrease in DNA methylation was observed alongside increased mRNA and protein abundance (**Figure 4C**). The remaining identified pathways presented a less clear relationship between OMIC layers.

### Part 2 – Exercise training modality-specific omics signatures and integration

#### Methylome associations with aerobic and resistance exercise training

To compare the effects of different exercise modalities on the skeletal muscle methylome, we conducted a meta-analysis of aerobic (6 studies, n=474) and resistance training cohorts (3 studies, n=127) (**Supplementary Table 2**). Aerobic training resulted in significant changes in 66,012 DMPs (FDR < 0.05), with 47% hypo-methylated and 53% hyper-methylated post-exercise training (**Figure 5A, Supplementary Table 17**). Specifically, 2,762 CpGs were located in active TSS and 12,390 in enhancer regions. The proportion of hypermethylation in these regions were similar to the observed for all CpGs, with 38% of CpGs at active TSS and 35% at enhancers being hypo-methylated. Additionally, we identified 8,479 DMRs (FDR < 0.05) associated with aerobic training, covering 8,752 genes.

**Figure 5.**
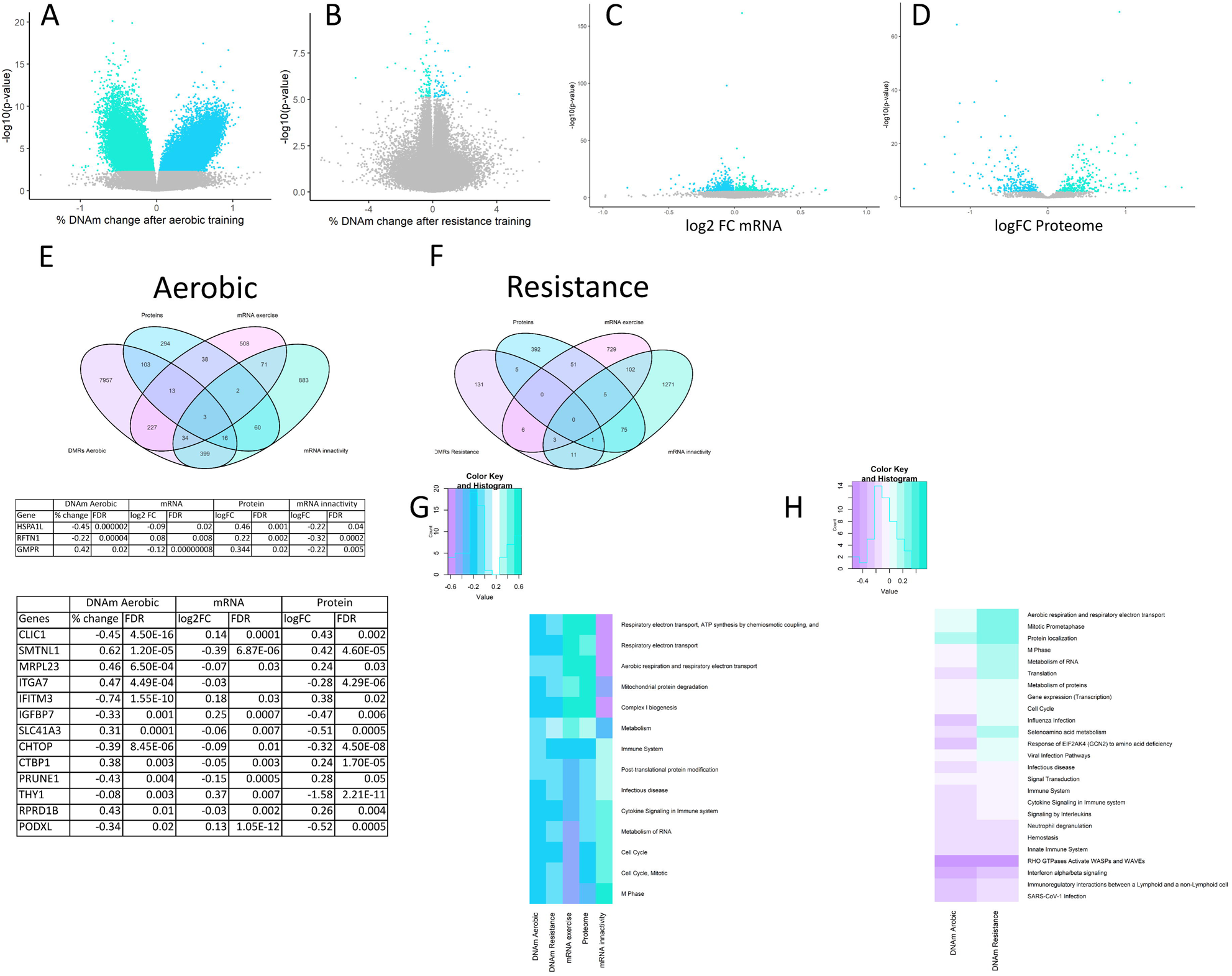
omics meta-analysis of distinct exercise modalities reveals a large effect on methylome after aerobic training but not after resistance training. **A)** Volcano plot of the effect size (% DNAm change after aerobic exercise training) and significance of exercise for each CpG included in the meta-analysis (pooled results of 9 independent muscle datasets); green dots represent differentially hypo-methylated CpGs, and blue dots represent differentially hyper-methylated CpGs at FDR <0.05. **B)** Volcano plot of the effect size (% DNAm change after resistance exercise training). **C)** Volcano plot of log2FC transcriptomic changes after exercise training. **D)** Volcano plot of log2FC proteomic changes after aerobic training. **E)** Venn diagram with intersected gene targets after aerobic exercise training and tables with intersected genes information. **F)** Venn diagram of intersected genes after resistance exercise. **G)** Top integration pathways representing a clear inverse relationship between exercise and inactivity. The blue colour represents an overall decrease in genes extracted from omics sites measured by the S distance. The band is represented in arbitrary units. **H)** Top DNA methylation integration pathways according to exercise training modality.

In contrast, resistance training resulted in minimal significant changes, with only 140 DMPs (FDR < 0.05), of which 61% were hypomethylated (**Figure 5B, Supplementary Table 18**). Post-resistance training, 19 CpGs were located in active TSS and 17 in enhancer regions. Interestingly, the direction of methylation changes differed in regulatory regions: 42% of CpGs in active TSS and 70% in enhancers were hypomethylated after resistance training. The DMPs from resistance training were annotated to 157 genes and did not form significant DMRs.

#### Transcriptome associations with aerobic and resistance exercise training

In the ExTraMeta database (Amar et al., 2021), comprising 952 samples, 896 mRNAs exhibited significant expression changes post-exercise training (FDR<0.05). 34% of these mRNAs showed increased expression levels (**Figure 5C & Supplementary Table 19**). Similarly, to the methylome analysis, the results revealed a predominant association with aerobic exercise, with 845 transcripts linked to it, whereas only 51 transcripts were associated with resistance training. Exercise-related mRNAs were enriched for 30 Reactome pathways (FDR <0.05 – **Supplementary Table 20**), with the most significant pathways related to mRNA splicing, muscle contraction, TCA cycle and respiratory electron transport.

Understanding the molecular responses to inactivity can provide insight into the mechanisms of muscle atrophy and understand muscle plasticity and adaptability. Thus, to provide a contrasting perspective, we also examined mRNA expression changes following a reduction in physical activity using a recent meta-analysis by Deane et al. (Deane et al., 2021). This analysis identified 1,468 mRNAs with altered expression following immobilization/disuse interventions, of which 46% were upregulated after immobilization (**Supplementary Table 21**). 110 transcripts overlapped between exercise and inactivity. Of these 72 (62 after aerobic and 10 after resistance) displayed an inverse relationship between exercise vs immobilization and 38 (34 after aerobic and 4 after resistance training) a direct relationship where expression either increased or decreased after both exercise training and immobilization.

#### Proteome associations with aerobic training

Our protein-wide association study identified 529 proteins with altered expression levels following exercise training (FDR < 0.05, **Supplementary Figure 5D** and **Supplementary Table 22**). Among these, 51% were upregulated. Unfortunately, we were unable to obtain any proteomics data for resistance training, precluding a modality-specific analysis for this OMICs discipline. The proteins related to exercise showed enrichment in 8 Reactome pathways, all of which were downregulated post-exercise. The most significant pathways were related to RNA metabolism and mRNA splicing (FDR < 0.05, **Supplementary Table 23**).

#### Methylome, Transcriptome and Proteome Integration

Following aerobic training, three genes, Heat Shock Protein Family A Hsp70 Member 1 Like (*HSPA1L*), Guanosine Monophosphate Reductase (*GMPR*), Raftlin, Lipid Raft Linker 1 (*RFTN1*), were common across all three OMIC layers from inactivity studies (**Figure 5E**). Interestingly, *RFTN1* showed decreased methylation, increased mRNA and protein levels post-exercise, and decreased mRNA levels post-immobilization. In contrast, *HSPA1L* exhibited decreased methylation and mRNA expression after exercise but increased protein expression and decreased mRNA expression after immobilization. *GMPR* displayed increased methylation after training, decreased mRNA expression after exercise as well as inactivity, yet an increase was observed at the protein level. Thirteen genes (Chloride Intracellular Channel 1 (*CLIC1*), Smoothelin-Like 1 (*SMTNL1*), Mitochondrial Ribosomal Protein L23 (*MRPL23*), Integrin Subunit Alpha 7 (*ITGA7*), Interferon Induced Transmembrane Protein 3 (*IFITM3*), Insulin Like Growth Factor Binding Protein 7 (*IGFBP7*), Solute Carrier Family 41 Member 3 (*SLC41A3*), Chromatin Target Of PRMT1 (*CHTOP*), C-Terminal Binding Protein 1 (*CTBP1*), Prune Exopolyphosphatase 1 (*PRUNE1*), Thy-1 Cell Surface Antigen (*THY1*), Regulation Of Nuclear Pre-MRNA Domain Containing 1B (*RPRD1B*), Podocalyxin Like (*PODXL*)) were common across DNAm, mRNA, and proteome data after aerobic training (**Figure 5E**). Conversely, post-resistance training, only one gene Microtubule-Associated Protein 4 (*MAP4*) was detected across DNAm, mRNA post-resistance training, and mRNA after immobilization (**Figure 5F**).

To explore the methylome in a modality-specific manner, we conducted a comprehensive gene set enrichment analysis (GSEA) to identify pathways common to both training modalities (Kaspi and Ziemann, 2020). We identified 316 Reactome pathways enriched for DNAm following both aerobic and resistance training (**Figure 5G** and **Supplementary Table 24**). Pathways associated with cancer suppression (e.g., CTNNB1 S33/S37/S45/T141 mutants, signalling by GSK3beta mutants), insulin, and thyroid regulation (e.g., IRS activation, thyroxine biosynthesis) showed increased enrichment across both training modalities. Conversely, pathways linked to infectious diseases and immunity were downregulated following both types of exercise (**Figure 5H**, **Supplementary Table 25**). A notable difference was observed in cell cycle-related pathways between aerobic and resistance training. DNAm associated with aerobic training appeared to decrease cell cycle pathways, whereas resistance training led to an increase (**Figure 5H**). Due to limitations in separating mRNA meta-regression by modality and the fact that our proteomics meta-analysis included only aerobic training samples, we could not conduct modality-specific pathway analysis for other OMICS layers.

## Discussion

We conducted an in-depth multi-OMIC analyses in human skeletal muscle to uncover molecular patterns, including TFs, associated with CRF and type of exercise training. We identified distinct molecular signatures and novel genes across the methylome, transcriptome, and proteome layers, associated with higher VO_2max_ levels. Additionally, we explored the modality-specific effects of aerobic (n=601) and resistance training (n=160) on DNA methylation, revealing significantly larger number of CpGs associated with aerobic training compared to resistance training. This disparity might be due to the larger sample size for aerobic training (n=601) compared to resistance training (n=160), as well as individual variability. However, it is also plausible that these differences reflect distinct molecular adaptations between exercise types and/or variability in training length, intensity, and volume (Egan and Sharples, 2023).

DNA methylation changes in response to exercise are transient but also cumulative (Barres et al., 2012), and thus the magnitude of changes at the expression level in trained muscle at rest may be smaller compared to untrained muscle responding to an acute exercise bout (Sexton et al., 2023; Turner et al., 2019). Such difference highlights the dynamic nature of DNA methylation, where gene expression changes from each acute exercise bout accumulate over time to produce long-term protein adaptations (Seaborne et al., 2018). Thus, we suggest that the large disparity in the number of significant CpGs between aerobic and resistance training identified in this study may reflect the methylation changes retained in trained individuals as a response to exercise, distinguishing them from those in untrained individuals. The inclusion of a wide range of training intensities (i.e. high-intensity, low-intensity) within the aerobic cohorts may capture a broad spectrum of methylation changes, partly explaining the significant differences observed between the results of aerobic and resistance training.

We identified 9,822 CpG sites strongly associated with baseline VO_2max_. The enrichment of hypomethylated CpGs with higher VO_2max_ aligns with previous studies linking hypomethylation to increased fitness levels (Plaza-Diaz et al., 2022; Seaborne et al., 2018a; Seaborne et al., 2018b; Sexton et al., 2023; Sharples et al., 2016). Our regional analysis unveiled novel pathways previously unreported in the context of CRF at the DNA methylation level. For instance, Antigen Processing: Ubiquitination & Proteasome Degradation pathway, primarily involved in immune response and protein turnover, and the NEDDylation pathway, a post-translational modification process crucial for protein regulation and signalling, represents novel findings with limited exploration in relation to CRF. Recent studies have shown that acute exercise increases NEDDylation-associated enzymes, highlighting the regulation of the ubiquitome at the proteome level (Parker et al., 2020). These proteins, conjugated with Cullin-RING E3 ligases, enhance ubiquitin activity during exercise. The enrichment of both pathways at the methylome level suggests that these regulatory mechanisms may respond to long-term exercise and are associated with higher CRF levels, indicating a crucial role for epigenetic regulation in adapting to exercise and improving fitness. Additional pathways related to immune response and signalling, that have not previously been associated with CRF, include TRAIL signalling, NFE2L2 regulating anti-oxidant/detoxification enzymes, and the activation of type I IFN production by LRR FLII-interacting Protein 1 (LRRFIP1). For instance, TRAIL signalling, which induces apoptosis, has been suggested as a mediator of muscle recovery, with post-exercise plasma analysis being associated with accelerated recovery (Jameson et al., 2022). The identification of this pathway at the methylome level in skeletal muscle suggests not only upstream regulation but also potential cross-talk between muscle and blood. However, it is important to consider that the observations in this study might reflect broader systemic changes since bulk biopsies were used without cell-type-specific deconvolution (Seaborne and Ochala, 2023). NFE2L2, a key regulator of the antioxidant response, has been extensively studied in animal models (Kitaoka et al., 2019; Wafi et al., 2019; Wang et al., 2019). There is also evidence suggesting that epigenetic downregulation may confer protection against osteoporosis (Chen et al., 2021a). Our study reveals a heightened presence of this pathway linked to increased CRF. Together with the aforementioned research, this underscores the interconnectedness of bone and muscle health.

We identified 162 transcripts and 351 proteins associated with baseline VO_2max_. Integrating the OMIC layers revealed dynamic relationships, where DNA methylation changes showed an inverse correlation with mRNA and protein changes in genes associated with mitochondrial function. Pathway analysis highlighted metabolic pathways, including the TCA cycle and respiratory electron transport, as significantly associated with VO_2max_ across all OMIC levels. These findings suggest that DNAm plays a critical role in modulating gene expression patterns related to mitochondrial metabolism in response to exercise (Li et al., 2022). Noteworthy, multiple pathways related to the cell cycle showed downregulation with increasing VO_2max_ across all OMIC levels. This observation aligns with the physiological demands of exercise, where muscle contraction, oxygen delivery, and metabolic processes require substantial energy expenditure (Hawley et al., 2014). As a result, the body may prioritise energy allocation towards supporting immediate exercise needs, potentially diverting resources from non-essential processes like cell division.

Interestingly, when examining the interaction between DNA methylation, TFs, and transcriptome results, we found that Paired Box 8 (*PAX8*) was targeted by five TFs. However, its expression was significantly regulated only by DNA methylation, and the direct effect of TFs on *PAX8* was not significant, indicating that the TFs role was not applicable. Furthermore, the lack of cell type specific analysis is a limiting factor, as we cannot untangle if such relationships are occurring in skeletal muscle cells, satellite cells or other interstitial cells within the tissue niche. Nonetheless, through the integration of multiple OMICs layers, we uncovered the consistent repression of cell cycle pathways across DNA methylation, mRNA, and protein levels. This finding underscores the intricate interplay between exercise-induced metabolic adaptations and cell cycle regulation, revealing a novel aspect of exercise-related molecular responses that extends beyond conventional mitochondrial-focused analyses.

Our analysis revealed various scenarios where gene expression levels could possibly be influenced by DNA methylation, TFs, or both. DNA methylation can primarily regulate gene expression, or it can jointly regulate it with TF activity. Additionally, TFs can regulate gene expression independently of DNA methylation, driving target gene expression solely through TF activity (Silva et al., 2022). We found 20 TFs that were enriched in hypo- or hyper-DMRs associated with VO_2max_. Significant transcripts associated with VO_2max_, were mostly located in genomic regions containing TFBS for enriched TFs identified in our DNA methylation analysis. This suggests an indirect feedback loop between DNA methylation and transcription. Epigenetic modifications, such as DNA methylation, can modulate the binding affinity of TFs, thereby influencing the transcriptional activity of genes associated with VO_2max_. While our data suggest that TFs may be the primary regulators of gene expression, surpassing the influence of CpGs associated with VO_2max_, which aligns with previous studies demonstrating the interplay between DNA methylation and transcriptional regulation (Lindholm et al., 2014). However, a limitation of this association is that UniBind accesses all tissue types, meaning these enrichments may not be specific to skeletal muscle; experimental processes are necessary to confirm these relationships. To further explore these interactions, we performed an analysis between DNA methylation, TFs, and the transcriptome using *MethReg*. Although this required a matched sample set and thus a smaller number of samples were available, we found that TFs in the exercise context generally act as activators for gene expression while the effect of DNA methylation on TFs was not significant, meaning that regardless of methylation levels being low or high methylation alone did not stop the function of TFs modulating gene expression. Even though TFs may appear to be activators in this context when looking at the interaction between TFs and DNA methylation we observed that most targets exhibited a significant synergistic effect. A limitation of Part 1 of our study is the heterogeneity in VO_2max_ assessment methods, which could introduce variability. Our study provides novel insights into the molecular mechanisms underlying exercise-induced gene regulation and adaptation, potentially uncovering new targets for enhancing exercise adaptation and health.

In our study five genes have shown congruence at all OMIC levels with VO_2max_, highlighting their significance in exercise responses. Although not all these genes have been studied in the context of exercise, most have been investigated for their roles in cellular stress response, and metabolism. ***NCEH1*** is involved in lipid metabolism, particularly in the hydrolysis of cholesterol esters. Reduced *NCEH1* levels in mice fed a high-fat diet led to impaired endothelium-dependent relaxation (EDR) in the heart, while increasing *NCEH1* expression restored normal EDR function (Sun et al., 2024) suggesting that *NCEH1* plays a critical role in maintaining cardiovascular health by regulating lipid metabolism and endothelial function. In our study, we observed a decrease in DNA methylation alongside increased expression at the transcriptome and proteome levels in individuals with higher VO_2max_, suggesting a significant influence of *NCEH1* in relation to CRF and underscoring its potential role in cardiovascular health and exercise adaptation. Our findings highlight *NCEH1* as a key player in the molecular mechanisms underlying the benefits of exercise, making it a potential target for therapeutic strategies aimed at improving cardiovascular health and fitness. ***ALDH6A1*** encodes a mitochondrial enzyme involved in the metabolism of amino acids and carbohydrates. Downregulation of *ALDH6A1* has been suggested as a marker of insulin resistance in type 2 diabetes (T2D) (Liu et al., 2022). Similar to *NCEH1*, our results indicate a deep regulation of *ALDH6A1* associated with higher CRF, which may confer a protective role against the development of T2D. ***HSPA2*** belongs to the heat shock protein family and is involved in cellular stress response and protein folding. Increased expression of *HSPA2* has been linked to sarcopenia (i.e. muscle wasting) in the Taiwanese population (Chen et al., 2021). Our findings showed a decrease in *HSPA2* expression with higher CRF, with all OMIC levels, including DNA methylation, being downregulated. The decrease in all OMIC levels observed with higher CRF likely reflects complex interplays involving epigenetic modifications, metabolic adjustments, and broader physiological responses. Further research is needed to understand the mechanisms behind these findings. ***PARK7*** encodes a protein involved in oxidative stress response and mitochondrial function. While initially studied in the context of Parkinson’s disease (Sanyal et al., 2020; Zhang et al., 2016), it has not been extensively reported in the context of exercise. However, *PARK7* has been shown to regulate *NRF2* (Zhang et al., 2021), which is part of the *NFE2L2* pathway, a target uncovered in our study. ***CAB39***, also known as MO25, is involved in regulating AMP-activated protein kinase (AMPK) signalling pathways (Liang et al., 2020). While no direct link between *CAB39* and exercise has been established, AMPK is a key regulator of cellular energy homeostasis, activated during exercise to enhance glucose uptake and fatty acid oxidation (Wu et al., 2024). By activating AMPK, *CAB39* likely mediates the cellular and molecular changes required for efficient energy production, improved endurance, and overall metabolic health. This positions *CAB39* as a key potential target for understanding how the body adapts to exercise and responds to metabolic stress, promoting increased CRF.

Exercise modality-specific analyses revealed 13 intersected genes (*CLIC1, SMTNL1, MRPL23, ITGA7, IFITM3, IGFBP7, SLC41A3, CHTOP, CTBP1, PRUNE1, THY1, RPRD1B,* and *PODXL*) associated with aerobic exercise across all OMIC layers, excluding mRNA followed by inactivity. In contrast, resistance training had only one intersecting gene (*MAP4*). Several of these genes (*SMTNL1, MRPL23, ITGA7, IGFBP7, SLC41A3, CHTOP, CTBP1, THY1* and *MAP4*) have previously been reported to be modulated by exercise (Chiang et al., 2019; Chou et al., 2023; Kilic-Erkek et al., 2021; Leckie et al., 2023; Lee et al., 2022; Liberman et al., 2022; Lueders et al., 2011; Luo et al., 2024; Murali and MacDonald, 2018; Wooldridge et al., 2008; Zou et al., 2011). However, five genes (*CLIC1, IFITM3, PRUNE1, RPRD1B* and *PODXL*) appear to be newly associated with aerobic exercise training. ***CLIC1*** acts as a sensor of oxidative stress, and its inhibition has been shown to protect against cellular senescence and endothelial dysfunction via the *NRF2* pathway (Lu et al., 2021). In our study, *CLIC1* expression increased, likely in response to the exercise-induced stress. The AMPK-mediated signalling pathway activates *NRF2* through phosphorylation and nuclear translocation, which is also regulated by *CLIC1* (Lu et al., 2021). Additionally, *CLIC1* activation contributes to H2O2-induced mitochondrial dysfunction and activation of mitochondrial fission. Overexpression of *CLIC1* has been reported to inhibit *NRF2* nuclear translocation. Our findings suggest that higher CRF is positively associated with *NRF2*, while aerobic training also increasing *CLIC1*. This indicates a temporary stress response leading to greater adaptation and a synergistic mechanism between *CLIC1, NRF2*, and AMPK signalling, making *CLIC1* a promising candidate for future studies on exercise adaptation (Lu et al., 2021). ***IFITM3*** is a defence gene, whose reduction leads to decreased CD8+ T cell levels in aged individuals after COVID infection (Hou et al., 2022), suggesting its crucial role in immune response during viral infections. Conversely, increased expression of *IFITM3* has been linked to normal ageing and Alzheimer’s disease (Hur et al., 2020), indicating its involvement in age-related immune regulation and neurodegenerative processes. Therefore, the significance of the increased *IFITM3* expression observed after exercise in our study remains unclear. ***PRUNE1*** is implicated in various cellular processes, including cell proliferation, mitigation and survival. Its expression has been shown to be positively correlated with tumour metastasis (Ferrucci et al., 2024). Although there is no direct evidence linking *PRUNE1* to exercise, its involvement in cellular processes such as cell proliferation, differentiation, and tissue repair can be indirectly associated with exercise. Exercise-induced muscle adaptation and repair processes could potentially be influenced by the mechanisms in which PRUNE1 is involved, highlighting its relevance in the context of exercise biology. ***RPRD1B*** is known to function in transcriptional regulation and RNA processing. It acts as a transcriptional co-regulator and is involved in the regulation of RNA polymerase II activity, which is crucial for the transcription of protein-coding genes (Ni et al., 2011). More research is needed to investigate the role of this gene in response to exercise, and to whether it is a pivotal regulator of transcription in exercise training. ***PODXL*** is involved in cell adhesion and migration. The expression of p53 is known to inhibit *PODXL* (Kim et al., 2007), however, its role in response to exercise remains unclear.

In summary, our large-scale multi-OMIC analyses of human skeletal muscle have uncovered previously unexplored genes and pathways related to CRF at baseline. We have identified significant overlaps between DNA methylation, enriched TFs and TFBS with associated transcriptome targets. Although some of our intersected genes have not been previously studies in the context of exercise we observed based on previous studies that genes associated with CRF appear linked to ageing-related diseases and exhibit an inverse pattern in individuals with higher CRF in our study. Our investigation into modality-specific responses to exercise training provide strong evidence of how CRF levels and adaptation to exercise are associated with multiple pathways across all three OMIC layers, despite limitations due a smaller number of studies on DNA methylation and proteomics after resistance training. Nonetheless, our results provide compelling evidence for the association of CRF levels and exercise adaptation with multiple pathways across all three omics layers, highlighting new potential targets for future research.

## Acknowledgments

We would like to express gratitude to all of our participants and our collaborators and their efforts during the intervention making this study possible. We would also like to thank Dr Sarah Voisin, a previous member of our lab, for the guide and advice, in particular in the bioinformatic analyses. This work was supported by Nir Eynon’s NHMRC Investigator Grant (APP1194159). The Gene SMART study is also supported by an Australian Research Council (ARC) Discovery Project Grants (DP190103081 & DP200101830). This study used BPA-enabled (Bioplatforms Australia)/NCRIS-enabled (National Collaborative Research Infrastructure Strategy) infrastructure located at the Monash Proteomics and Metabolomics Facility. Study PXD023084 was supported by supported by an unconditional donation from the Novo Nordisk Foundation (NNF) to NNF Center for Basic Metabolic Research (http://www.cbmr.ku.dk) (Grant number NNF18CC0034900), NNF Center for Protein Research (https://www.cpr.ku.dk/) (Grant number NNF14CC001), and Team Denmark. KHP: Research council of Finland (#335443, 314383, 272376, 266286), Finnish Medical Foundation, Gyllenberg Foundation, Novo Nordisk Foundation (#NNF20OC0060547, NNF17OC0027232, NNF10OC1013354), Finnish Diabetes Research Foundation, Paulo Foundation, University of Helsinki and Helsinki University Hospital, Government Research Funds. MO: Research council of Finland (#328685, 307339, 297908 and 251316), Sigrid Juselius Foundation, Liv O Hälsa r.f., and Minerva Foundation. Open access publishing facilitated by Monash University, as part of the Wiley - Monash University agreement via the Council of Australian University Librarians.

## Author contributions

MJ and NE, conceptualized and led the project, contributing to all aspects of development and writing. Collaborators including SL, AG, APS, RS, JS, AH, ES, MO, JB, NZ, OH, OE, OA, SL, CS, CB, MW, IL, AET, LG, IB, RAES, JWH, NRH, LMH, LRG, KJA, AD, and BJF contributed to data collection and manuscript revision.

## Methods

### Participants

We used multiple cohorts for each OMIC layer (**Figure 1** & **Supplementary Tables 1 & 2).** For easy reference we have subdivided our method in two parts. Part 1 will detail studies and methods included in the cross-sectional analysis between OMICs layers and CRF. Part 2 will detail studies and methods included in the OMICs analysis of exercise training. Note that some studies were included in both part 1 and 2, in which case they will not be described twice but rather reference to description above will be pointed out. A summary table for studies included in each part as well as statistical models including covariates for each study is found in supplementary tables 1 and 2, for part 1 and part 2 respectively.

#### Part 1 – Cross-sectional meta-analysis of OMICs and its associations with CRF

<u>Study cohorts:</u>

<u>The full information on study cohorts including sample size, age, sex, phenotype/s, and type of exercise intervention can be found in Supplementary Tables 1 and 2.</u>

<u>Statistical Analysis used in both Part 1 & 2</u>

### DNA methylation pre-processing and analyses

DNA methylation data was pre-processed using the *ChAMP* analysis pipeline (Morris et al., 2014; Tian et al., 2017) and *minfi* packages in the R statistical software version 4.0.2. First, we removed probes located on the sex chromosomes. Both males and females were analysed and pre-processed together. Then, we removed probes with detection p-value > 0.01, with < 3 beads in at least 5% of samples, with missing β-values, aligning to multiple locations, and non-CpG probes. Probes mapping to single-nucleotide polymorphisms (SNPs) or located close to SNPs showing a high frequency in the Caucasian population (“EUR” population probes described in Zhou *et al*. (Zhou et al., 2017)) were also removed. β-values were then obtained as follows:

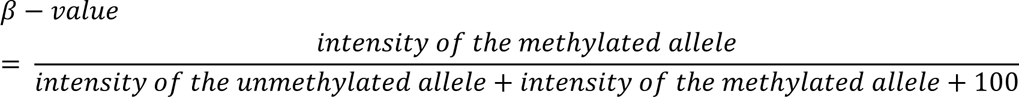

Then, we applied β-mixture quantile normalisation (BMIQ) to normalize the distributions of Type I and Type II probes. We then performed singular value decomposition (SVD) to identify major sources of variability in the dataset. This data exploration step revealed that batch and position on the batch were significant sources of variation in the data. Therefore, data was converted to M-values according to the following formula:

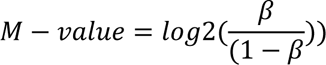

and the ComBat function of the *sva* package (Leek et al., 2012) was used to remove those technical artefacts.

For DNA methylation analyses we first converted β-values to M-values as M-values are more homoscedastic and therefore more appropriate for differential analysis of methylation levels (Du et al., 2010). We used linear models and moderated Bayesian statistics as implemented in *limma* (Ritchie et al., 2015). DNA methylation levels at each individual CpG were regressed against CRF for part 1 and timepoint for part 2. To avoid confounding effects and reduce influence of external factors to the analysis other covariates were included for each individual study and details are found in **Supplementary Tables 1 & 2**. When necessary we used the duplicateCorrelation function to account for the paired design (i.e. repeated measures on the same individuals or twin data). Results obtained from each individual study were then meta-analysed using an inverse variance weighted meta-analysis with METAL (Willer et al., 2010). CpGs associated with *CRF* and *Timepoint* at a false discovery rate (FDR) < 0.05 were considered differentially methylated positions (DMPs). If DMPs were identified, we then proceeded to identify differentially methylated regions (DMRs), i.e. contiguous clusters of DMPs showing consistent changes in DNA methylation. DMRs were identified using the *DMRcate* package (Peters et al., 2015). We then proceeded to identify pathways potentially altered as a result of the differential DNA methylation. We performed a gene set enrichment analysis to identify enriched pathways using the Reactome database and *mitch* (Kaspi and Ziemann, 2020). All pathways at FDR <0.05 were considered significant. Finally, we conducted chromatin states enrichment analyses in our DMRs detailed explanation has been previously reported here (Voisin et al., 2021).

### Transcriptomics analyses

Results from the independent transcriptomics datasets were combined using an inverse variance weighted meta-analysis with METAL (Willer et al., 2010). We used METAL since it does not require all datasets to include every mRNA site in all studies. Different sets of mRNAs may be filtered out during pre-processing of each individual dataset, which means the overlap between the datasets is imperfect and a given mRNA may only be present in 1 out of 4 datasets, or 2 out of 4 datasets. For robustness, we only included mRNAs present in 2 out of the 4 cohorts. We used a fixed effect (as opposed to random effect) meta-analysis, assuming one true effect size of exercise on the transcriptome, which is shared by all the included studies. Nevertheless, Cochran’s Q-test for heterogeneity was performed to test whether effect sizes were homogeneous between studies (a heterogeneity index (I^2^) >50% reflects heterogeneity between studies). The mRNAs associated with fitness at a stringent meta-analysis FDR <0.05 were deemed significant.

### Proteomics pre-processing and analyses

Before normalisation, proteomic data was filtered for high-confidence protein observations. In addition, contaminants, proteins that have been only identified by a single peptide and proteins not identified/quantified consistently across the experiment have been removed. The remaining missing values were imputed using the missing-not-at-random (MNAR) method, assuming the missingness was due to low expression for such proteins, which are then normalised using variance scaling normalization (VSN). Both imputations and VSN were conducted by the *DEP* package (Zhang et al., 2018). Batch effects were normalised with the internal referencing scaling (IRS) method (Plubell et al., 2017) by the use of reference channels. We used linear models and moderated Bayesian statistics as implemented in *limma* (Ritchie et al., 2015). Results from each study were then meta-analysed using METAL (Willer et al., 2010). Results with an FDR <0.05 were deemed significant. Pathway analyses was conducted using the Reactome and *mitch* (Kaspi and Ziemann, 2020).

#### Integration of methylome, transcriptome and proteome profile according to CRF and changes after exercise intervention

Integration analyses was conducted by using a multi-contrast pathway enrichment analyses for multi-omics data (mitch) and is described in detail elsewhere (Kaspi and Ziemann, 2020). Hypergeometric testing is used to calculate the probability of obtaining a given number of successes in a sample drawn from a finite population without replacement. This test is particularly useful in genomics for identifying whether a subset of genes (e.g., differentially methylated genes) is significantly enriched in a predefined set of genes (e.g., a biological pathway or gene set). We used this method to confirm our interactions were not simply due to chance using the *missMethyl* package (Phipson et al., 2016).

We used the ggplot2 (Ginestet, 2011), ggpubr (Kassambara, 2018) packages for data visualisation.

#### Integration of methylome, TFs and transcriptome

TFs and TFBS were derived from significant CpGs associated with CRF using UniBind tool (Puig et al., 2021). UniBind provides a comprehensive database of TFBS derived from ChIP-seq data, helping to identify potential regulatory elements involved in gene expression. We also investigated the interaction between DNA methylation and TFs using matched samples from the Gene SMART cohort, which included 89 samples. For this analysis, we employed the *methReg* tool (Silva et al., 2022), which integrates DNA methylation data with gene expression and TF binding information to uncover regulatory relationships.

